# Yttrium-iron garnet film magnetometer for magnetic microparticles in vivo registration studies

**DOI:** 10.1101/2022.12.29.522185

**Authors:** N. Koshev, P. Kapralov, S. Evstigneeva, O. Lutsenko, P. Shilina, M. Zharkov, N. Pyataev, A. Darwish, A. Timin, M. Ostras, I. Radchenko, G. Sukhorukov, P. Vetoshko

**Author notes:** Corresponding author: N.Koshev.

## Abstract

In the current article, we present a new kind of magnetometer for quantitative determination of magnetic objects in biological fluids and tissues. The sensor is based on yttrium-iron garnet film with optical signal registration system. Inheriting the working principle of a fluxgate magnetometers, the sensor works at a room-temperature, its wide dynamic range allows the measurements in an unshielded environment. A small size of sensitive element combined with a short recovery time after the excitation coils are off provide us with a potentially high spatial and temporal resolution of measurements. We show the feasibility of the sensor by sensing the remanent magnetization of Magnetic Nanoparticles (MNPs) both in vitro (test tubes, dry MNPs) and in vivo (local injection of the MNPs into mice).

## 1 Introduction

### 1.1 Nanoparticles magnetic relaxometry and imaging

Nowadays, various visualization techniques based on the magnetic signals registration in living organisms are of great importance in medicine. These methods can be divided into two groups: 1) Magnetic Resonance Imaging (MRI) 2) Various types of magnetometry. MRI based on the effect of resonant absorption of electromagnetic radiation by hydrogen nuclei placed in a strong magnetic field. MRI is one of the most informative methods for diagnosing neoplasms [1], as well as different vascular and dystrophic diseases [34] [11] Despite several undoubted advantages, MRI has certain disadvantages, including the technical complexity and bulkiness of the equipment [35]. It significantly reduces the availability of this method.

A magnetometry can be considered as alternative to MRI, which is based on the signals registration (relaxation time and/or remanent magnetization) from magnetic contrast agents introduced into the circulatory system of a living organism. As a rule, these agents are super-paramagnetic or ferromagnetic nano- or microparticles. Their magnetic moments distributed over a volume under investigation are oriented along the applied saturating field with enough magnitude. After the saturating field is off, the MNPs’ magnetic moments starting to re-orient with time, which leads to dependence of the net magnetic moment on time (relaxation curves). The single domain particles are subject to two main magnetic relaxation mechanisms: Neel and Brownian relaxations [17]. The first one depends mostly on the properties of MNPs, while the second is affected by the properties of the area, which contains MNPs. The Magnetic Relaxometry (MRX) method is based on measurement of the dependence of the net magnetic moment on time in rapidly decaying saturating magnetic field. The registered signal depends on applied field magnitude, on spatial distribution of MNPs over the volume of interest, and on the relative positions of sensor. Evaluation of the magnetic field induction on the surface and inside the living organism makes it possible to reconstruct the distribution of MNPs in the whole volume of the organism, which reflects the accumulation in certain organs and tissues, as well as to evaluate the intensity of blood flow. The totality of the obtained data can allow to detect tumor nodes, as well as areas of tissue ischemia.

While the MRX method is mostly devoted to investigation of the magnetic relaxation at some point, the Magnetic Particle Imaging (MPI) method aims at reconstruction of the pointwise concentration of particles distributed over the object of interest volume [50]. MPI is based on receiving signal from the non-linear re-magnetization response of SPIONs to an oscillating magnetic field [18] [9]. Also, highly sensitive systems include MPI scanner, operating on a two-frequency method. This is a powerful radiation-free method, giving the spatial resolution as of 1mm. Despite current MRI’s spatial resolution is higher (25 100μm), the submillimeter spatial resolution of the MPI approach can be obtained with improvement of sensitivity of the sensors. Among the advantages of the MPI method over the MRI/CT we can highlight radiation-free nature, much more compact devices, lower capital and operational expenses.

From the previous discussion one can see that sensitivity is the key factor for both MPI and MRX methods. This issue is related to high-sensitive magnetometers also being essential for the current research.

#### 1.1.1 High-sensitive magnetic sensors

Conventionally, the most common high-sensitive magnetometers are presented with Superconducting Quantum Interference Devices (SQUID). These magnetometers utilize the phenomenon of Josephson currents interference in a superconductive conditions. The SQUIDs show high sensitivity (up to 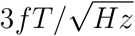) and stability, but have several significant drawbacks which makes their usage non-reasonable in some cases. SQUIDs demand cooling with liquid helium (see e.g [2] [38, 43] or liquid nitrogen [3, 4, 16] in order to maintain superconductivity phenomenon. Due to this, SQUID-based devices are quite bulky. The sensor arrays are not flexible or adaptable, and, due to cooling, needed to be placed at a thick walls Dewar containers, which decrease the resulting sensitivity while increase the distance between sensor and object under investigation. The main drawback of SQUID-based MRX is related to comparatively long recovery of SQUIDs after the pumping magnetic field is off. This significantly limits the relaxation times, allowing to research only the Brownian relaxation and the remanent magnetization. Despite all above, the SQUID-based measurements of MNPs still can provide us all required information due to variety of experimental techniques. The research [23]shows the change of magnetic properties of MNPs in dependence on properties of media (solution). The properties were explored by changing DC fields in a measurement of 1s, which is much greater than both Neel and Brownian relaxation times. The concentrations of MNPs were 50 μg/ml. A very promising SQUID-based in-vivo experiment was shown in [30], where the sensitivity of the experimental setup was enough to resolve nanoparticles with the mass of 1.2mg Fe injected in the rabbit.

In MPI invivo experiments authors [5] detected 1.1*ηg* of iron (SNR 3.9) with the spatial resolution about 600 *μm,* but the excitation field was 45 kHz, 20 mT peak amplitude, that are not available for clinical use, especially with living organisms. On the laboratory version scanner in [19] signals from 5ng were received in vitro and in vivo experiments on mice 500ng were shown and the flow of an intravenously injected indicator through the mouse heart was recorded. The main disadvantage of such systems is high-frequency magnetic fields usage, which can cause electromagnetic stimulation in living organisms.

Recent scientific and technological progress results in the development of the new types of high-sensitive sensors. Among them, atomic optically-pumped magnetometer (OPM) is one of the most promising high sensitive magnetometer. Compared to SQUID, the OPMs are a bit less sensitive (up to 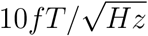), but significantly outperform SQUIDs in terms of compactness. These sensors do not demand the liquid helium and can be arranged in flexible sensor arrays. All these advantages lead to dramatic increase of popularity of them during the last decade. OPMs, however, also subject to some drawbacks related to instability and long calibration procedure, which depends on the temperature. The lasers of OPMs are subject to degradation with time, and heating make these magnetometers demand solutions for heat removal. However, there are several research for overcoming these disadvantages. For example, research [31, 40] demonstrate non-zero field sensors, which theoretically might resolve the question of stability and re-calibration, while [**?**,29] shows the possibility of development of He4 based room temperature OPMs, resolving the heating problem. The approach is very promising despite the sensitivity of the latter is currently below the ’common’ Rb OPMs.

In terms of MRX/MPI, we need to highlight the OPM-MRX research [26], [25]. The first research allowed to register an iron concentration of 5.8 *μg* with application of Quspin QZFM OPMs; the second work [25] demonstrates a brilliant results on usage of OPMs based on free-precession principle. The gradient metric system working in an unshielded environment showed the sensitivity of 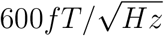, and the 1.37 *μg* iron have been detected.

Another type of sensors feasible for MRX is a fluxgate one. These sensors have a wide dynamic range and very low heat dissipation, which also make them suitable for unshielded measurements. The research [32] shows the relaxometry measurements with the fluxgate sensors with intrinsic noise of 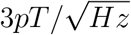, which allowed to register MNPs with 5.6μg of iron. The same research team publishes in 2022 work [24],where the experimental setup with sensitivity up to 254 *ng* of iron claimed and proven in experiment.

The publications above show the advantages of some types of OPMs and fluxgate sensors over SQUIDs in terms of MRX and MPI. First, both types of magnetometers show lesser recovery times, which allow to measure relaxometry curves as opposed to SQUIDs. Despite the fact that remanent can be measured with SQUIDs, such measurements may be laborious due to specifics of SQUIDs measurements during the excitation [6]. Currently, the main advantage of SQUIDs over OPMs and fluxgate sensors is related to higher sensitivity. However, the situation tends to change during the last decades. While the OPMs are approaching in terms of sensitivity to SQUIDs, recent advantages in fluxgate technologies also show that these sensors can reach much better sensitivity level. E.g., in the work [27] we presented a first flux-gate sensor with sensitivity a bit lower but of the same order with OPMs 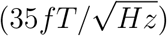. This sensor type works at room temperature, does not need any cooling or heating, subject to wide dynamic range. The same type sensors but with optical system of regestration of signal was described in [48].This sensor registers a change of the polarization angle in garnet films. This sensor can detected magnetic field about 10pT (the magnetization field 50kHz).In terms of relaxometry, the main advantage of the flux-gate sensors is related to quite short recovery time after the pumping coils are off power.

In this article we show new type of high-sensitive magnetometer for MRX analysis. Based on the yttrium-iron garnet films (YIG), the sensor in general inherits the working principle of the fluxgate magnetometers but differs in terms of reading the film magnetic state. In this study, only the results of the analysis of the remanent magnetization of particles are presented.The research is devoted to the feasibility study and is organized as follows. In the section (II) we describe magnetometer, experimental setup, and experimental paradigm, and MNP used for our research. The section III is devoted to the representation of our results in terms of noise study, in-vitro study (test-tubes tests with MNPs), and primary in-vivo study, where we register the remanent magnetization of MNPs injected in mice. There is also another type of sensors based on the GMI effect. In the study [15] a ferromagnetic soft magnetic microconductor CoFeSiBNb was used as a sensitive element of the magnetometer. It was possible to register only a sample of Fe3O4 400 micrograms by weight of iron magnetite (2.8 *mg/ml*). Signals from samples with a lower mass of iron, which were originally stated in the article, were not given. The sensitivity of the magnetometer according to the data given is not large enough to detect biologically applicable masses.

## 2 Methods and Materials

### 2.1 Experimental setup

The experimental setup includes the magnetometer, excitation coils, and control system based on data acquisition unit NI USB-2650.

#### 2.1.1 Description of the sensor

The sensitive element of the sensor is the thin yttrium-iron garnet (YIG) film grown on GGG base. The film was chemically etched to the round shape with the radius of about 15 mm. Applying the constant magnetic field of magnitude 3 Oe, we can form the single-domain magnetic state in the film. The magnetic state of the film is sensed using the magneto-optic Faraday effect. The ray of the solid-state laser with the wavelength 532 *vm* coming through the Glan-Taylor prism and falls on the YIG film with the angle closed to the Brewster’s angle (see Fig. 1, a). The Glan-Taylor prism is being used to eliminate the polarization noises of the laser. After coming through the YIG film, the light being separated to two orthogonal polarization components using the Wollaston prism. The difference in light intensity in these two components is being measured using the balanced optosensor, which allows to avoid influence of the laser intensity noise at output. The photo of the experimental setup is presented on Fig. 1, b. We further refer the magnetometer as FYIG.

**Figure 1:**
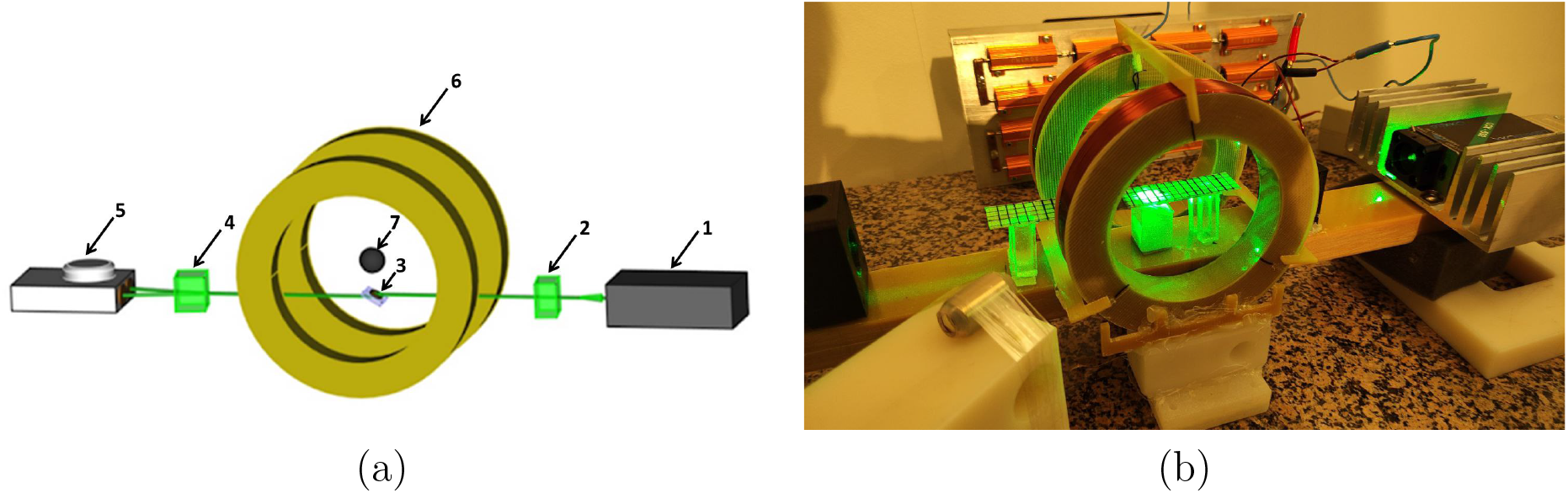
MO-YIG based experimental setup. (a) sheme FYIG: 1 - the solid-state laser, 2 - the Glan-Taylor prism, 3 - YIG film, 4 - the Wollaston prism, 5 - the balanced optosensor, 6 - the impulse coils, 7 - the measured sample; (b) photo of the setup

#### 2.1.2 Excitation system

The magnetization system consists of Helmholtz coils, which create a pulsed magnetic field, and a constant magnetic field source, which is necessary to saturate the magnetic film FYIG (much less than the pulsed one). A current was applied to the Helmholtz coils for a time of 20 ms, which made it possible to create an approximately uniform magnetic field of 12 mT to magnetize the nanoparticles. Next, the pulsed magnetization field is turned off, and during the next 80 ms the magnetic field created by the demagnetized nanoparticles is measured. After that, current is again supplied to the coils, but of opposite polarity, and the measurement cycle is repeated. Bipolar magnetization is used to minimize the effect of irreversible particles magnetization. Polarity switching is performed by a specially designed switch, which is resistant to self-induction voltage surges up to 1000V. This made it possible to dissipate the energy of the magnetic field of the impulse coils in less than 1 ms.

#### 2.1.3 Materials

Since in vivo magnetometry is currently at the development stage, there are many unresolved problems also from the side of the contrast agents development. The main requirements for them are biocompatibility (low toxicity) and high remanence (high signal for registration). Potentially, these requirements are met by MNPs based on iron oxide and one of the objectives of this work was their contrasting ability evaluation. Several types of magnetic particles were used for research: submicron spherical particles Fe3O4, coated with sodium polystyrene sulfonate (Fe3O4@PSS/nanosphere), submicron Fe3O4 rods coated with sodium polyacrylate (Fe3O4@PAA/rods). Magneto electro nanoparticles (MENs) offer a special combination of crucial properties to address significant obstacles in contemporary cancer treatment, in contrast of classic nanomaterials, MENs show unique characterization because of the existence of a non-zero ferroelectricity effect, given their large surface area, core-shell magnetoelectric nanoparticles with ferromagnetic cores which thus demonstrate magnetostriction and ferroelectric shells that demonstrate piezoelectricity have the capability to generate mV voltage energy at the nanometer range [8, 42]. These nanoparticles differ from classical ones such as in their high capacity for image-guided targeted therapy, which can be used for targeting framework guided by physical force to improve the cell viability of bioactive molecules only across the carcinoma biological membranes, while sparing normal ones, with an external monitoring process to discharge drugs on demand throughout an external influence of the electric fields that underpinning the intrinsic reaction mechanism between particular cells and the packed cargo due to the ferromagnetic and ferroelectric effect [37, 46]. Therefore, CoFe2O4 particles were also studied in pure form (CoFe2O4/no_shell) and in the piezoelectric shell of BaTiO3 (CoFe2O4@BaTiO3/thin_shell, thick_shell). We should note, however, that we had only a single amount of these particles equal to 5mg; thus, in the Sec. III we show only a remanent for this single concentration.

Morphology of synthesized MNPs was evaluated with the transmission electron microscopy (TEM) using a FEI Tecnai Osiris (USA) microscope with an operating voltage of 200 kV equipped with a SuperX EDS system for ultrafast element mapping. The samples were diluted with deionized water to an acceptable concentration and placed onto a special substrate (copper mesh coated with carbon) in the form of a solution and then dried for 30 minutes at ambient conditions. Fe3O4@PSS particles are spherical, monodisperse, with an average size about 200-300 nm. It is clearly noticeable that the larger spheres of Fe3O4@PSS have a rough appearance and consist of many small nanoparticles (Fig. 2, a). This indicates the self-assembly of nanoparticles and the formation of spherical structures. Many spheres have open pores. According to the TEM study, the Fe3O4@PAA nanorods are sufficiently monodisperse with an average length of 700 nm and a diameter of 30 nm (Fig. 2, b). The magnetic properties of the obtained nanospheres and rods were evaluated using an EZ11 vibration magnetometer (Microsense Inc., Lowell, USA) at room temperature (25 oC) in dried form. Figure 3 shows the magnetization curves of the obtained MNPs, which indicate their ferrimagnetic properties, while the remanent magnetization was 21 and 8.5 emu/g for Fe3O4@PSS and Fe3O4@PAA, respectively.

**Figure 2:**
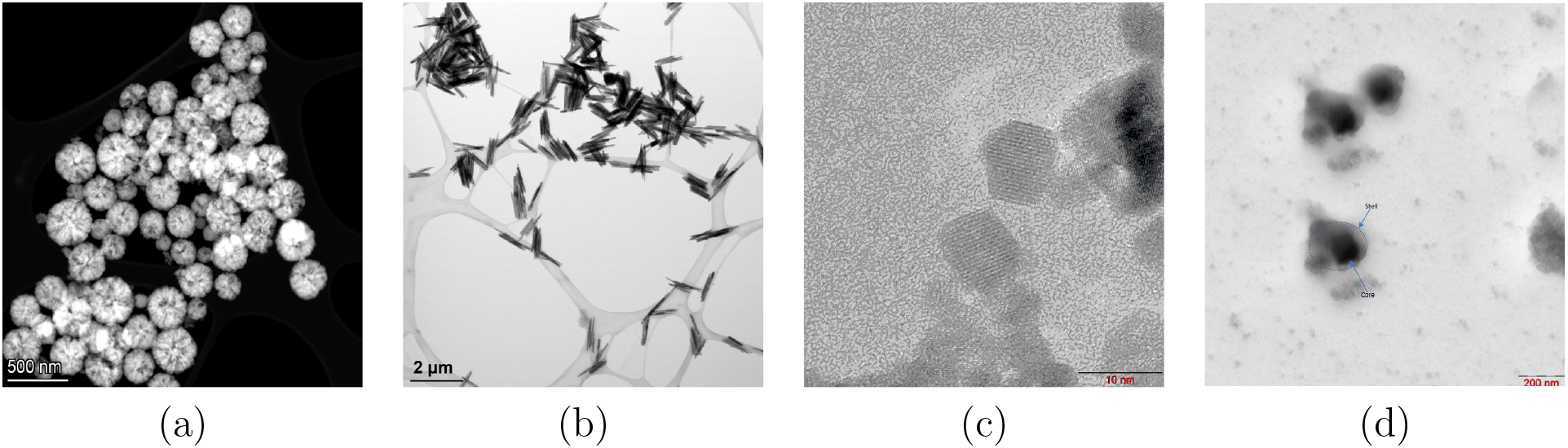
TEM images of investigated particles (a) - submicron spherical particles (Fe3O4@PSS); (b) - submicron rods (Fe3O4@PAA); (c) - particles CoFe2O4; (d) - particles CoFe2O4 in the piezoelectric shell of BaTiO3 (CoFe2O4@BaTiO3).

**Figure 3:**
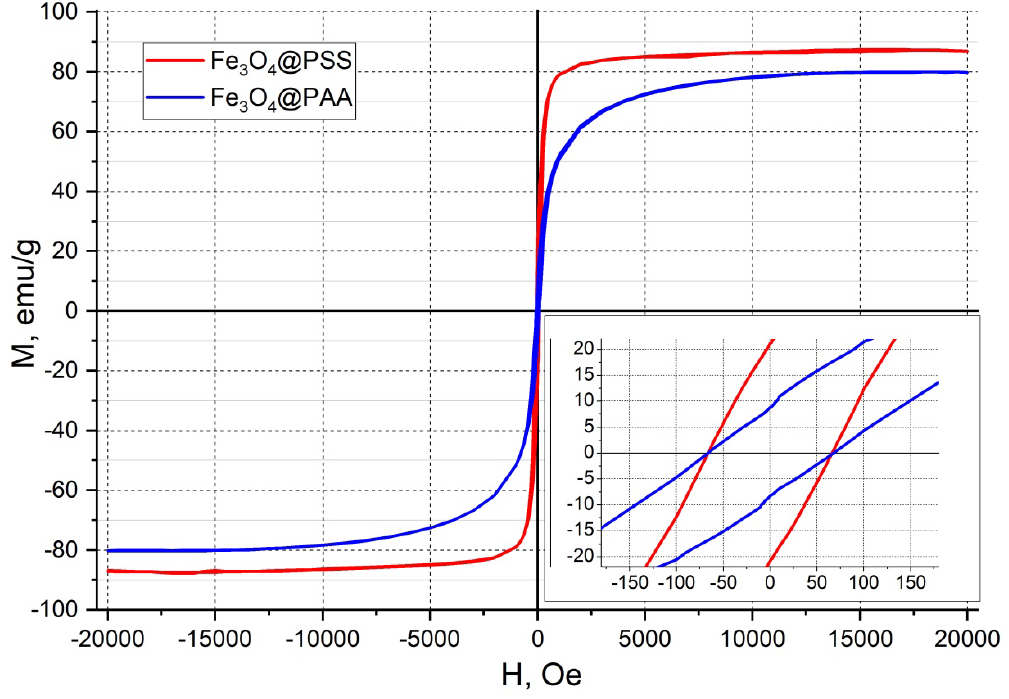
Magnetization curves for dry ’nanospheres’ (red curve) and ’rods’ (blue curve) samples.

CoFe2O4 MNPs were poly crystalline, and its crystal structure was of the inverse cubic spinel type. The TEM image of the CoFe2O4 sample is shown in Fig. 2, c. The image clearly shows that co-precipitation was used to create uniformly distributed, relatively homogenous cubic particles and the common particle size was 25 nm. The particle size of CoFe2O4@BaTiO3 were about 30nm (thin shell MENs) and 40 nm (thick shell MENs). Fig. 2, d shows a typical TEM image of the MENs with the core-shell structures clearly discernible.

### 2.2 Signal processing

In our experimental setup, the volume of interest is placed between the Helmholtz coils in the area of uniform saturating magnetic field. Under these conditions, we assume the magnetic moment at every point of the volume, containing the MNPs is oriented along this field. Denoting by Ω ⊂ ℝ^3^ the volume under investigation, we can write the impact to the B-field at sensor position **x**_*k*_ ∈ ℝ^3^ of the volume element with center of mass at the point *ξ* ∈ Ω as follows:

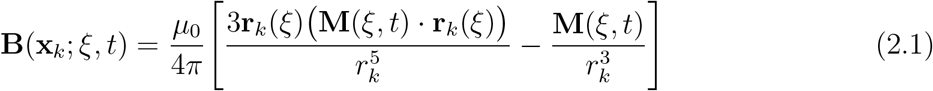

where **r**_*k*_ = *ξ* – **x**_*k*_, and *r_k_* = |**r**_*k*_|. The **M**(*ξ,t*) is the net magnetic moment of the volume element associated with the point *ξ* ∈ Ω at some moment of time *t*.

If the MNPs are immobilized (’dry’ experiments), the dependence of the magnetic moment **M** on time is governed only with the Néel relaxation model. If the particles are saluted in liquid, both Néel and Brownian mechanisms are presented. For the sake of generality we consider the last case in the current paragraph. When the excitation coils are off, then, after some time needed to cancel fully the saturation field, the net magnetic moment of the volume element *dV* associated with the point *ξ* is governed by the following model:

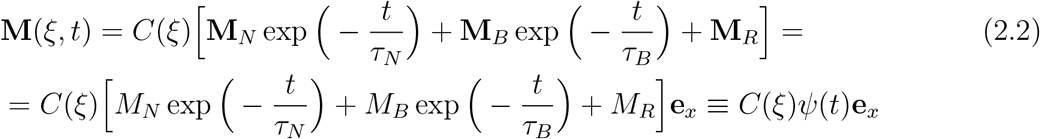

where *C*(*ξ*), ∈ Ω is the concentration of MNPs at the volume element *dV*, **e**_*x*_ is the unit vector along the X-axis, which we can orient along the excitation coils axis without loss of generality, and *M_R_* represents the remanent magnetization.

The whole B-field impact of the volume containing MNPs at sensor position can be obtained by integration of the above presentation over the whole domain Ω and takes therefore the following form:

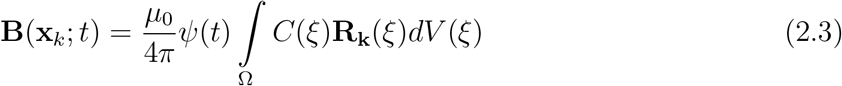

Here

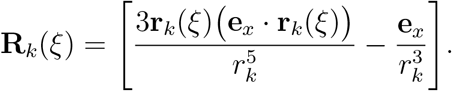

#### 2.2.1 Single sensor analysis

Analyzing the representation for the B-field (2.3), we can see that the measurements proportional to the function *ψ*(*t*). The function is decaying with time up to the constant, referred to be a remanent magnetization; the latter, obviously, depends on the measurement time. Analysis of the measurements for *t* ≤ *τ_N_* or *t* ≤ *τ_B_* means the MRX method of analysis Néel of Brownian relaxations respectively. However, in this case we need to reduce sufficiently the coils magnetic relaxation or, at least, subtract the coils relaxation from the registered relaxation curve. On the other hand, the magnitudes of the remanent are commonly lower than the magnetization at *t* < *τ_B_*. Thus, in the current article, to show the feasibility, we pay attention to the remanent magnetization, assuming the sensor is also suitable to the MRX due to short (about 0.5 ms) recovery time after the excitation coils are off.

The concept of remanent magnetization is related to the observation time [17]. In our experiments, the measurement time for the remanent magnetization analysis is *t* ∈ [40, 80] ms after the excitation coils are off. Since some of our experiments were provided in an unshielded environment, the measurements are subject to ’slow’ magnetic contamination, both periodic and non-periodic, which can impact at measured signal as constants or as slowly (with respect to the measurement time) changing functions. The latter contamination affecting the mean values of the voltage.

To separate these ’slow’ noises we use the statistical analysis of a number or measurements *B_i_*(*t*), *i* = 1, 2,…, *N_c_* ∈ [20, 500] registered within every experiment for the time *t* ∈ [40, 80] ms after the coils are off. The remanent magnetization 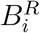 were then computed by averaging of every of registered curves. Since the distribution of these values subject to errors, we compute its statistical mean value and standard deviations. This approach, as opposed to conventional averaging of all measured curves, has several advantages. It allows to filter out the single values which are located farther from the mean than the standard deviation, avoiding impact of them in the resulting signal. Secondly, the statistical approach gives us an error of measurements in the meaning of the standard deviation, which might be crucial for the reconstruction of MNPs concentration distribution over a volume under investigation.

Exploring the remanent magnetization of low amounts of MNPs, we compare measurements of each experiment with the background noise also registered during these experiments.

Signal processing was done using Python and its modules Pandas, Scipy, and Numpy ([22, 39, 49]). The times needed for registration and processing of different number or curves are given below, in the section 3.

## 3 Results

### 3.1 Calibration and direction diagram

The calibration of the sensor has been done by application of the calibration coil with analytically calculated B-field at its center of 2*ηT* at the frequency of 980 Hz. The Fig. 4 a) shows the spectrum of calibration coil measurements. The sensitivity of the sensor is estimated to be 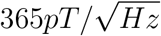 at 40 Hz. The spectrum allows us also to estimate the high-frequency intrinsic noise of the sensor of a level about 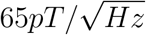.

**Figure 4:**
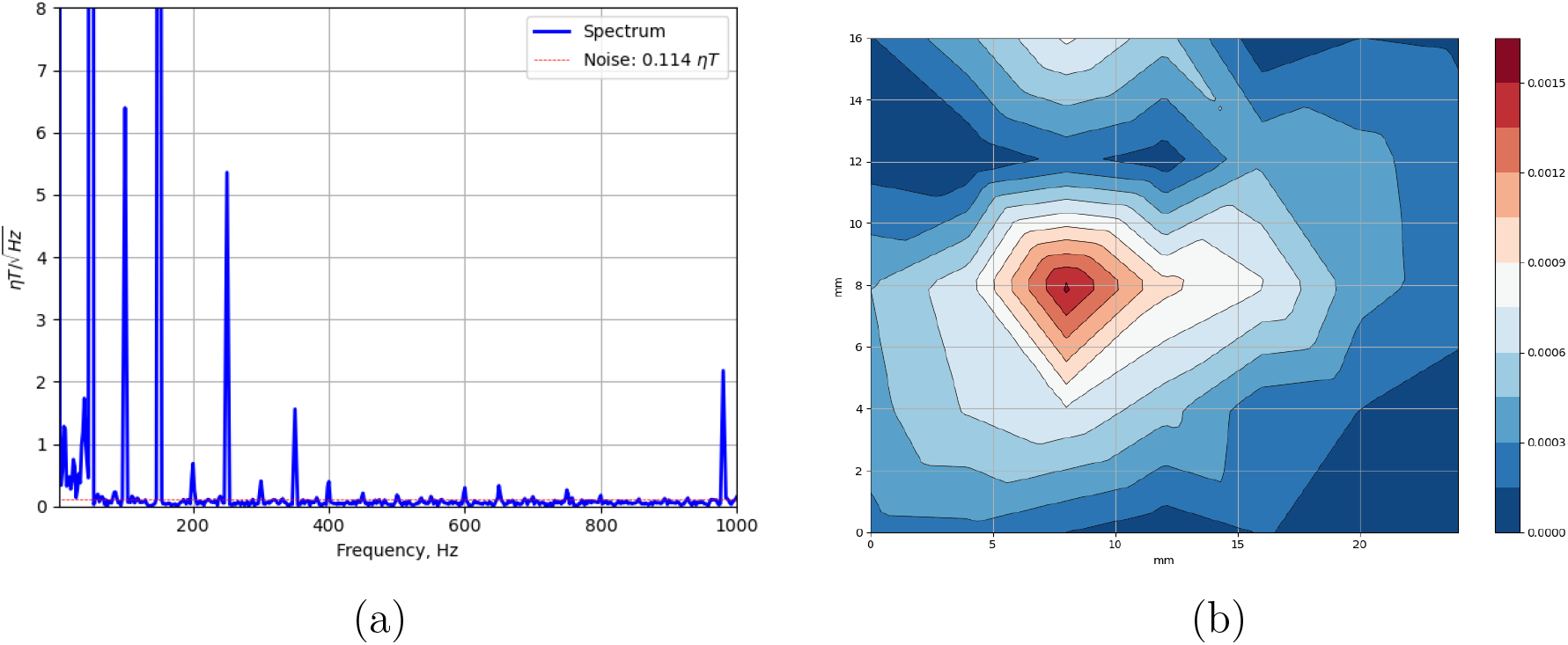
(a) Spectrum of the signal with calibration coil working at frequency of 980 Hz. (b) Direction diagram

The direction diagram (Fig. 4 b) shows the dependence of the signal amplitude on the position of a ’point’ source. The point source is simulated with a small coil working at frequency of 980 Hz and generating the B-field of 10*μT* at its center. The B-field is directed along the coil’s axis to simulate the MNPs field.

### 3.2 In-vitro study

In the in-vitro study, we investigated the signal generated by test tubes The test tubes study aims to explore the remanent magnetization of different samples of nanoparticles placed inside the test tubes. In the current experiments we use the ’dry’ samples described in the Table 1 with different concentrations. The results are shown in Fig. 5.

**Figure 5:**
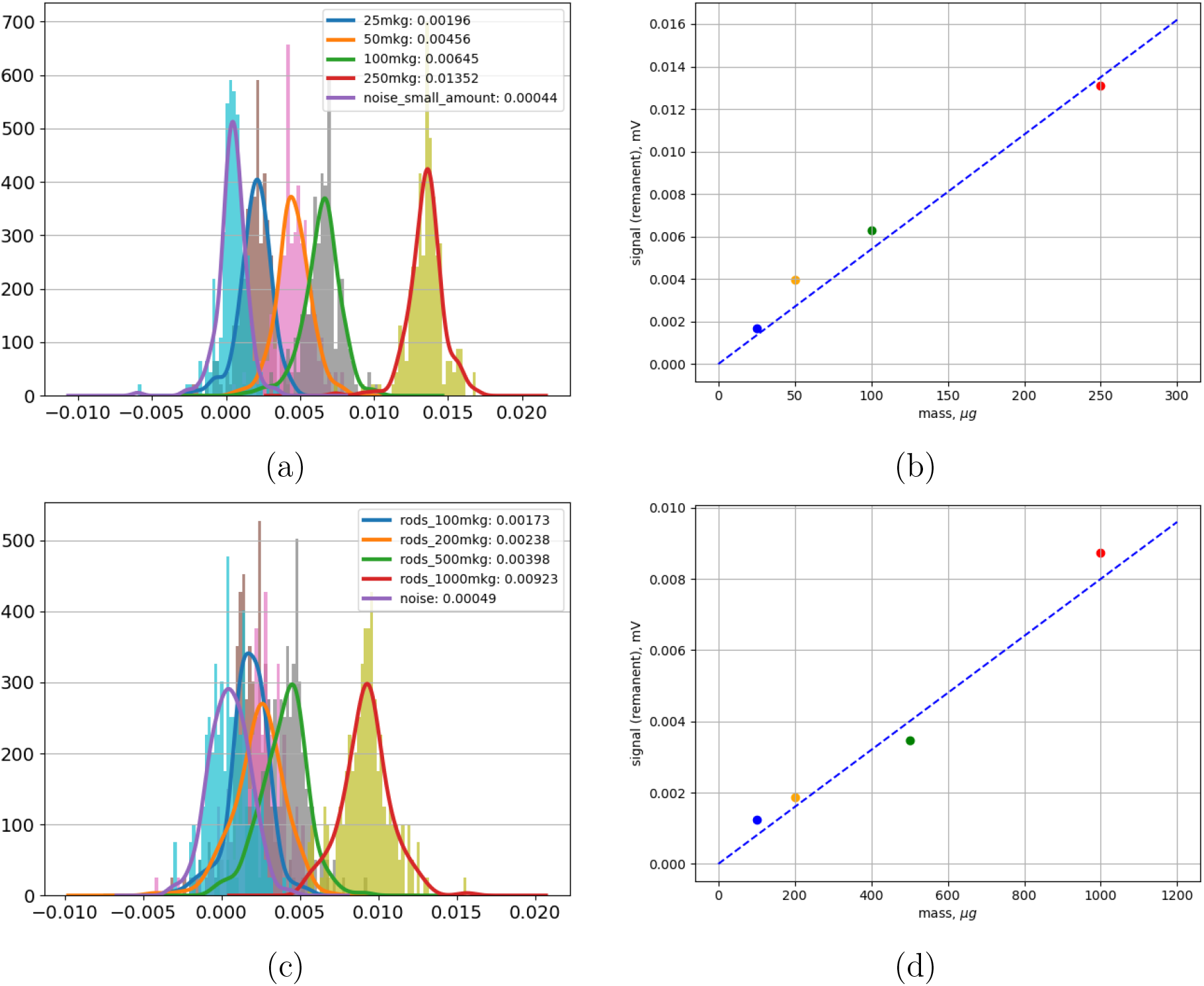
Remanent magnetization: (a) and (b) - histograms of measured signal, 200 measurements for every amount of ’nanospheres’ sample for 25, 50, 100, 250*μg*, and dependence of the mean signal on the amount; (c) and (d) the same for ’Rods’ sample, concentrations - 100, 200, 500 and 1000 *μg*.

**Table 1:** Description of different types of MNPs samples used for in vitro experiments

| Name | Material | Size, nm | Sensitivity, mV/mg |
| --- | --- | --- | --- |
| nanosphere | $Fe_3O_4@PSS$ | 200 – 300 | 0.05 |
| rods | $Fe_3O_4@PAA$ | 30/700 | 0.008 |
| no_shell | $CoFe_2O_4$ | 25 | 0.017 |
| thin_shell | $CoFe_2O_4@BaTiO_3$ | 25/3 | 0.005 |
| thick_shell | $CoFe_2O_4@BaTiO_3$ | 25/8 | 0.001 |

During our study, we were able to confidently observe the remanent magnetization for the 25 *μg* of ’nanospheres’ sample, and 100 *μg* of ’rods’ sample (Fig. 5, a). The CoFe2O4 and CoFe2O4@BaTiO3 MNPs were measured in only a single amount of 5 mg. All the measurements were conducted in unshielded environments, which explains rather high level of noise. The mean remanent B-field at sensor position can be obtained by subtraction of the mean noise value from the mean measurements. In order to validate our measurements, we studied also several other amounts of both samples (see Fig. 5, b)

The Fig. 6 shows the histogram of measurements of 3 samples for bio-compatible particles.

**Figure 6:**
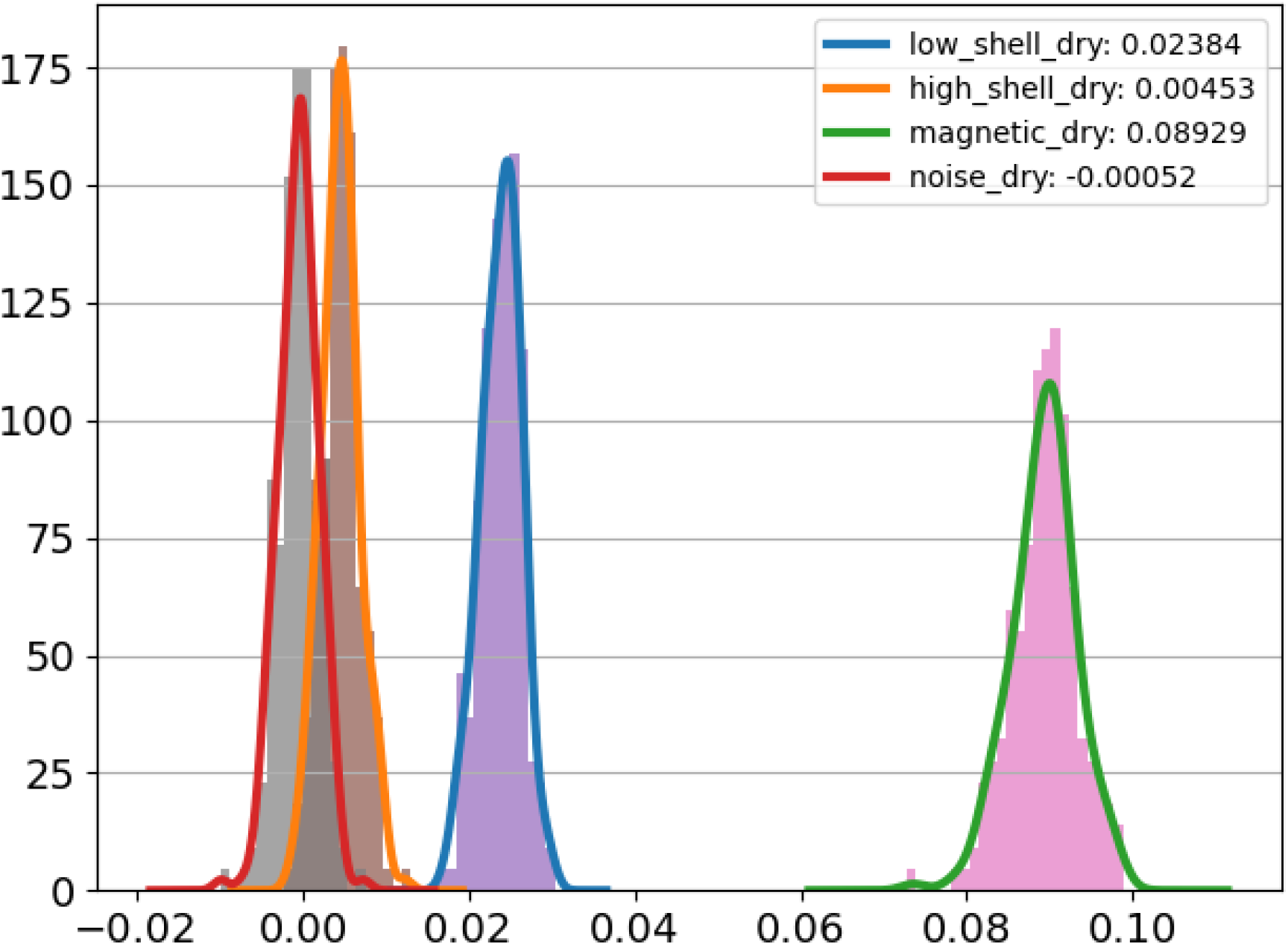
Histogram of measured remanent for bio-compatible particles: **name of samples** (200 measurements for every sample).

### 3.3 In-vivo experiments with mice

#### 3.3.1 Preparation

The experiment was conducted with male balb/c mice weighted at 20-25 g. Conditions of the experiments satisfied the Helsinki Declaration on Animal Rights and approved by the local committee on Ethics at National Research Mordovia State University. Animals were anesthetized with intra-abdominal injection of Zoletil (25 mg/kg) and Xylazine (5 mg/kg). The volume of the injected suspension was 0.1 ml. Magnetic substances were injected subcutaneously into the left lumbar region. The mass of nanoparticles was equal to 20, 10 and 5 mg of ’nanospheres’ sample, and 1 mg of ’rods’ sample.

#### 3.3.2 Measurements

Measurements: In the current research, we did not have the 3D scans of mice. For visual reference, we use the open source 3D model of the mouse and adapt its sizes and spatial placement to the exact coordinates of the measurement points and sensors. Thus, despite the 3D model is approximate, the coordinates of all points in the further figures are exact.

The measurement points defined a-priory on the skin of mice are shown in Fig. 7 (a). Since our experimental setup is single-sensor, we move and rotate mice in order to measure the magnetic field at the least distance to the measurement points. Applying the movements and rotations not to the mouse, but to the sensor, we form the array of sensor coordinates in a coordinate system associated with mice. The sensor positions obtained in such a way are depicted with Fig. 7 (b).

**Figure 7:**
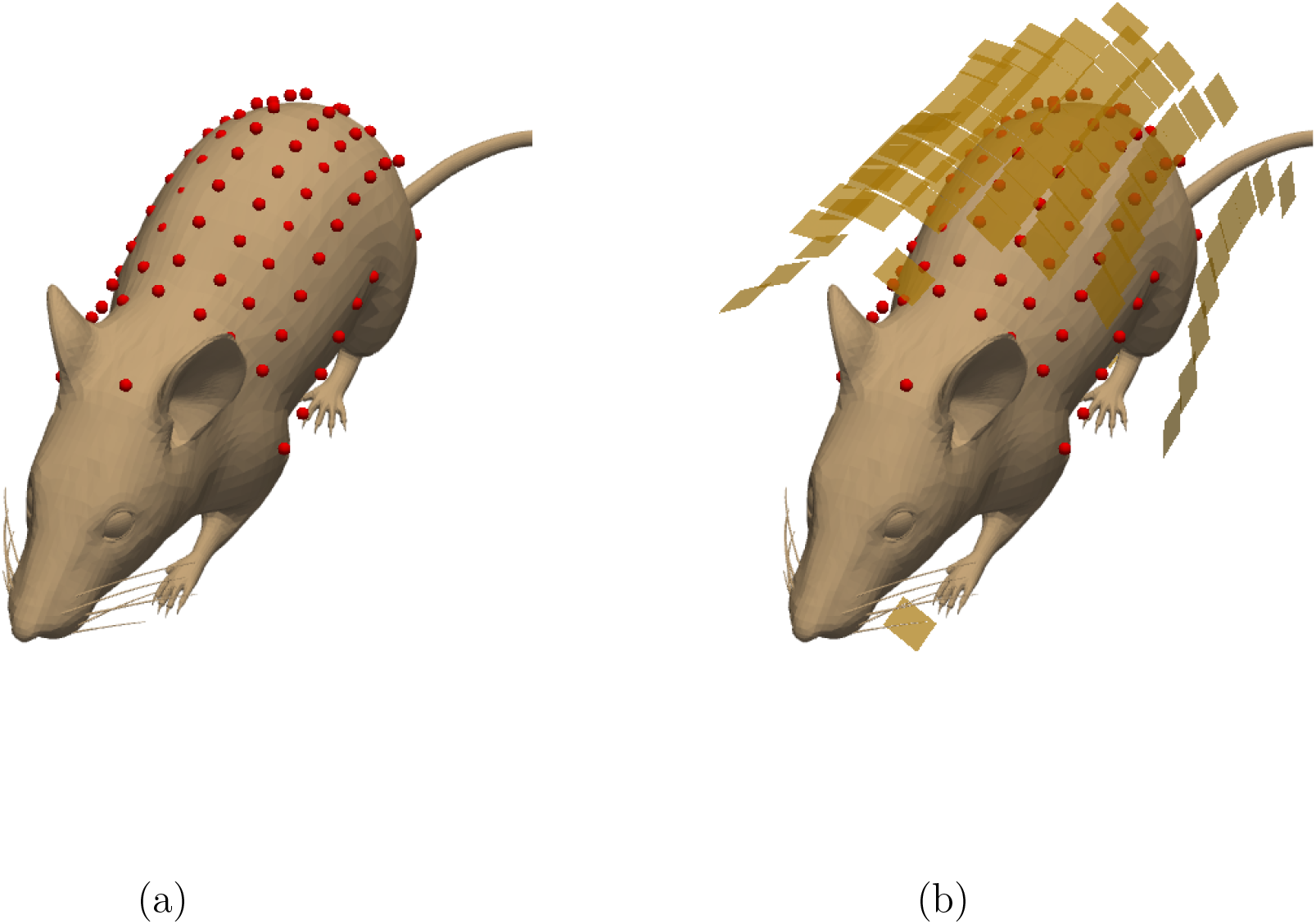
(a) - measurement points on the skin of mice (defined a-priory); (b) - sensors’ positions obtained via application of rotation and movement operations.

The Fig. 8 presents the measurements of the magnetic field at sensor’s positions. As mentioned earlier, the 3D model of mouse is shown for visual reference, but both sensors and injection positions are accurate. Due to the absence of the 3D model of the mouse, we do not apply any reconstruction techniques in the current article, leaving it for the further research

**Figure 8:**
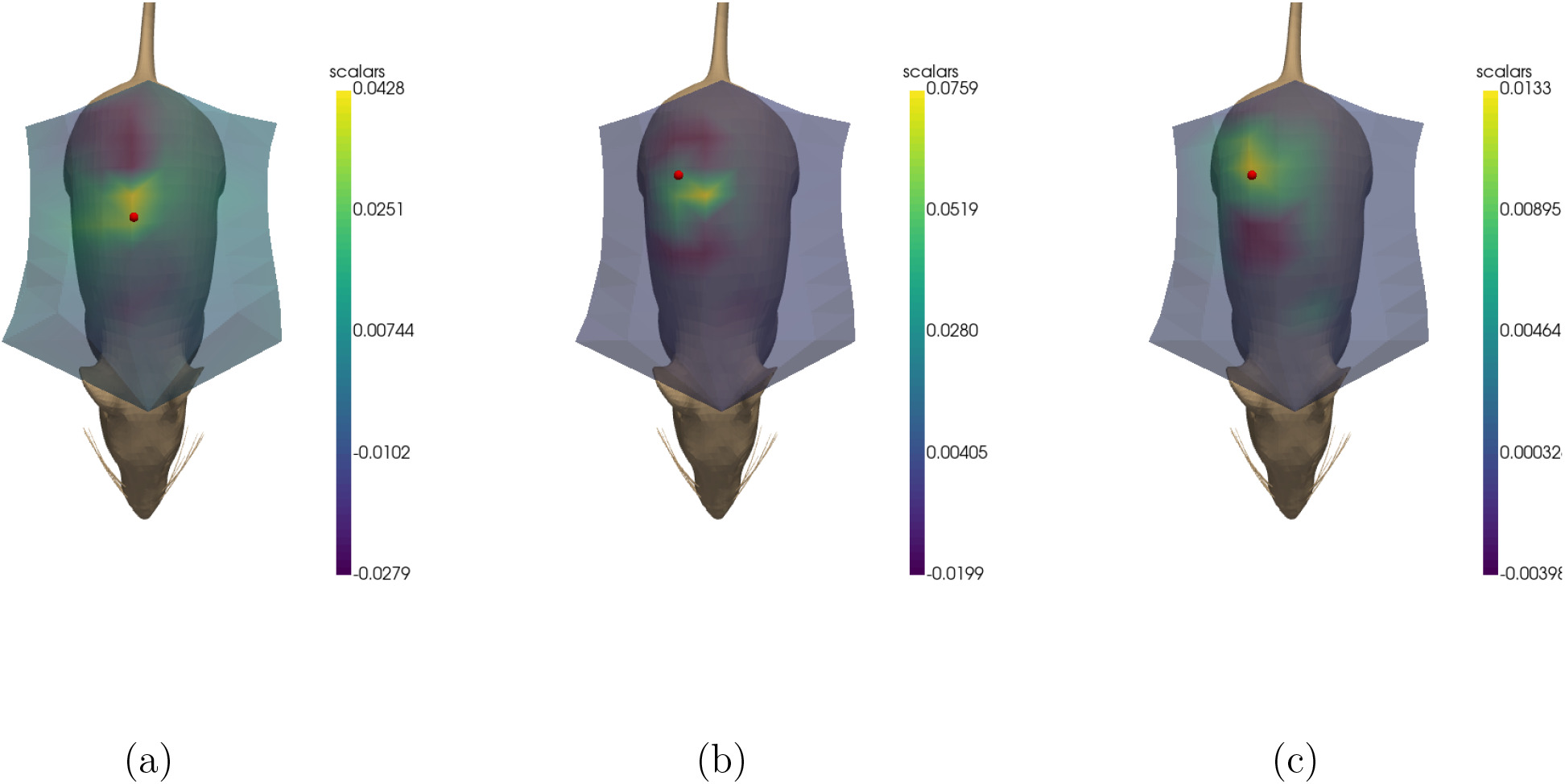
Measurements of the remanent magnetic field of samples, injected in mice. The red sphere means the injection point. (a) - 10 mg of the ’nanosphere’ sample; (b) - 5 mg of the ’rods’ sample; (c) - 1 mg of the ’rods’ sample.

## 4 Discussion

In the current research, we demonstrated a new FYIG sensor for MRX measurements. We used both in vitro and in vivo measurements of several types of MNPs showing the remanent due to comparatively big size, which avoids the superparamagnetic state. For two kinds, the measurements have been validated with dependence of the signal on the concentrations. The processing has been done within a statistical approach, which allowed us to register remanent for 25 *μg* nanospheres of *Fe*_3_*O*_4_ in the in-vitro tube test. In mice, the minimal amount we were using was 1 mg of the ’rods’ sample injected. Fig. 8, (c) shows that we can confidently register this amount of rods, which allows us to assume that lesser amounts will also be visible. The initial research presented in the current article provides us with several questions, issues, and ideas. The magnetic relaxation of MNPs will be the main topic of one of the future investigations. The remanent magnetization is much lower when compared to the magnetization at low times during the relaxation. Moreover, the sensor demonstrates quite short recovery times (about 0.5*ms*) after the excitation coils are off. Thus, we expect no serious difficulties in the application of this sensor to the magnetic relaxation. We need, however, to improve the excitation coils since the current ones subject to its own comparatively long (10 – 15 ms) relaxation after the power is off, which made impossible to investigate the MNPs magnetic relaxation.

One more serious aspect to investigate in the nearest future is the sensitivity of the sensor. Currently, the FYIG demonstrates sensitivity of about 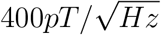. The sensor is based on the Faraday effect - the magnetization of the YIG film is being registered with a laser. The laser, however, also generates the noise. We expect that new generation of the YIG sensor will inherit one more of our sensors (see [27]), which will sufficiently improve the magnetometer in terms of sensitivity and make it more compact.

For purposes of magnetic particle imaging, we plan to develop mathematical and algorithmic base for inverse problem solution. The equation (2.3) allows us to state both forward and inverse problems of magnetic particle imaging and magneto relaxometry.While the forward modeling aims to estimate the signal at sensor knowing distribution of MNPs over the volume under investigation, the inverse problem, as opposed, dedicated to reconstruction of mass distribution of MNPs on the base of MRX/remanent measurements. Mathematically, both problems are closed to ones researched in the neuroimaging area (see e.g. [14, 20, 44]).

The forward modeling may be used to investigate different sensor arrays for effective MNPs imaging. Again, we plan to start our research on this with consideration of similar algorithms related to the neuroimaging area (see e.g. [21, 45, 47, 51]).

Being ill-posed by J.Hadamard, the inverse problem demands the development of new algorithms taking into account the specificity of the signal, noises, limitations of MNPs distributions. Some solutions were obtained for the MPI area [6,30]. A big number of comparatively effective methods were developed in the neuroimaging area. Recently, several promising approaches based on the solution of the ill-posed Cauchy problems for elliptic PDEs were proposed by various mathematicians [7, 10, 12, 13, 28, 33, 36], including some ML/AI-based methods [41]. One of the further works in the nearest future will be devoted to development of high-accuracy algorithms for reconstruction of the mass MNPs distribution and validation of it in the real MPI/MRX experiments.

## 4.0.1 Acknowledgements

The research related to synthesis of CoFe2O4 and CoFe2O4@BaTiO3 NPs was conducted under the financial support of the Ministry of science and higher education of Russian Federation FSEG-2022-0012.

## References

[1] A review on brain tumor diagnosis from mri images: Practical implications, key achievements, and lessons learned. Magnetic Resonance Imaging, 61:300–318, 2019.

[2] Natalie L Adolphi, Dale L Huber, Howard C Bryant, Todd C Monson, Danielle L Fegan, JitKang Lim, Jason E Trujillo, Trace E Tessier, Debbie M Lovato, Kimberly S Butler, et al. Characterization of single-core magnetite nanoparticles for magnetic imaging by squid relaxometry. Physics in Medicine & Biology, 55(19):5985, 2010.

[3] Lau M Andersen, Robert Oostenveld, Christoph Pfeiffer, Silvia Ruffieux, Veikko Jousmäki, Matti Hämäläinen, Justin F Schneiderman, and Daniel Lundqvist. Similarities and differences between on-scalp and conventional in-helmet magnetoencephalography recordings. PLoS One, 12(7):e0178602, 2017.

[4] Lau M Andersen, Christoph Pfeiffer, Silvia Ruffieux, Bushra Riaz, Dag Winkler, Justin F Schneiderman, and Daniel Lundqvist. On-scalp meg squids are sensitive to early somatosensory activity unseen by conventional meg. NeuroImage, 221:117157, 2020.

[5] Hamed Arami, Eric Teeman, Alyssa Troksa, Haydin Bradshaw, Katayoun Saatchi, Asahi Tomitaka, Sanjiv Sam Gambhir, Urs O. Häfeli, Denny Liggitt, and Kannan M. Krishnan. Tomographic magnetic particle imaging of cancer targeted nanoparticles. Nanoscale, 9:18723–18730, 2017.

[6] Daniel Baumgarten, Mario Liehr, Frank Wiekhorst, Uwe Steinhoff, Peter Münster, Peter Miethe, Lutz Trahms, and Jens Haueisen. Magnetic nanoparticle imaging by means of minimum norm estimates from remanence measurements. Medical & biological engineering & computing, 46(12):1177–1185, 2008.

[7] Frederik Berntsson and Lars Eld ’en. Numerical solution of a cauchy problem for the laplace equation. Inverse Problems, 17:839–853, 2001.

[8] Soutik Betal, Moumita Dutta, LF Cotica, A Bhalla, and R Guo. Batio3 coated cofe2o4–core-shell magnetoelectric nanoparticles (csmen) characterization. Integrated ferroelectrics, 166(1):225–231, 2015.

[9] Caroline Billings, Mitchell Langley, Gavin Warrington, Farzin Mashali, and Jacqueline Anne Johnson. Magnetic particle imaging: Current and future applications, magnetic nanoparticle synthesis methods and safety measures. International Journal of Molecular Sciences, 22(14):7651, Jul 2021.

[10] L. Bourgeois. Convergence rates for the quasi-reversibility method to solve the cauchy problem for laplaceâs equation. Inverse Problems, 22:413–430, 2006.

[11] Jedrzej Burakiewicz, Christopher DJ Sinclair, Dirk Fischer, Glenn A Walter, Hermien E Kan, and Kieren G Hollingsworth. Quantifying fat replacement of muscle by quantitative mri in muscular dystrophy. Journal of neurology, 264(10):2053–2067, 2017.

[12] Hui Cao, Michael V. Klibanov, and Sergei V. Pereverzev. A carleman estimate and the balancing principle in the quasi-reversibility method for solving the cauchy problem for the laplace equation. Inverse Problems, 25:25pp, 2009.

[13] Hui Cao and Sergei V. Pereverzev. The balancing principle for the regularization of elliptic cauchy problems. Inverse Problems, 23:19431961, 2007.

[14] J.C. De Munck, C.H. Wolters, and M. Clerc. EEG and MEG: forward modeling, chapter Chapter 6, pages 203–272. CUP/BRRR, 2012.

[15] Basile Dufay, Sébastien Saez, Matthieu Denoual, Christophe Dolabdjian, F Ludwig, E Heim, M Schilling, L Melo, A Yelon, and D Menard. Magnetorelaxometry of nanoparticles using a gmi magnetometer. Sensor Letters, 7(3):429–432, 2009.

[16] MI Faley, J Dammers, YV Maslennikov, JF Schneiderman, D Winkler, VP Koshelets, NJ Shah, and RE Dunin-Borkowski. High-tc squid biomagnetometers. Superconductor science and technology, 30(8):083001, 2017.

[17] Raluca Maria Fratila and Jesús Martínez De La Fuente. Nanomaterials for magnetic and optical hyperthermia applications. Elsevier, 2018.

[18] Bernhard Gleich and Jürgen Weizenecker. Tomographic imaging using the nonlinear response of magnetic particles. Nature, 435(7046):1214–1217, 2005.

[19] Matthias Graeser, Tobias Knopp, Patryk Szwargulski, Thomas Friedrich, Anselm von Gladiss, Michael Kaul, Kannan M Krishnan, Harald Ittrich, Gerhard Adam, and Thorsten M Buzug. Towards picogram detection of superparamagnetic iron-oxide particles using a gradiometric receive coil. Scientific reports, 7(1):1–13, 2017.

[20] M. Hamalainen, R. Hari, R. J. Ilmoniemi, J. Knuutila, and O. V. Lounasmaa. Magnetoencephalography: theory, instrumentation, and applications to noninvasive studies of the working human brain. Rev. Modern Phys., 1993.

[21] Matti S Hamalainen and Jukka Sarvas. Realistic conductivity geometry model of the human head for interpretation of neuromagnetic data. IEEE transactions on biomedical engineering, 36(2):165–171, 1989.

[22] Charles R. Harris, K. Jarrod Millman, Stéfan J. van der Walt, Ralf Gommers, Pauli Virtanen, David Cournapeau, Eric Wieser, Julian Taylor, Sebastian Berg, Nathaniel J. Smith, Robert Kern, Matti Picus, Stephan Hoyer, Marten H. van Kerkwijk, Matthew Brett, Allan Haldane, Jaime Fernández del Río, Mark Wiebe, Pearu Peterson, Pierre Gérard-Marchant, Kevin Sheppard, Tyler Reddy, Warren Weckesser, Hameer Abbasi, Christoph Gohlke, and Travis E. Oliphant. Array programming with NumPy. Nature, 585(7825):357–362, September 2020.

[23] Ryota Isshiki, Yuta Nakamura, Shun Takeuchi, Tetsuro Hirata, Kenji Sakai, Toshihiko Kiwa, and Keiji Tsukada. Evaluation of the magnetization properties of magnetic nanoparticles in serum using hts-squid. IEEE Transactions on Applied Superconductivity, 28(4):1–5, 2018.

[24] Klaas-Julian Janssen, Meinhard Schilling, Frank Ludwig, and Jing Zhong. Single harmonic-based narrowband magnetic particle imaging. Measurement Science and Technology, 33(9):095405, 2022.

[25] Aaron Jaufenthaler, Thomas Kornack, Victor Lebedev, Mark E Limes, Rainer Körber, Maik Liebl, and Daniel Baumgarten. Pulsed optically pumped magnetometers: Addressing dead time and bandwidth for the unshielded magnetorelaxometry of magnetic nanoparticles. Sensors, 21(4):1212, 2021.

[26] Aaron Jaufenthaler, Peter Schier, Thomas Middelmann, Maik Liebl, Frank Wiekhorst, and Daniel Baumgarten. Quantitative 2d magnetorelaxometry imaging of magnetic nanoparticles using optically pumped magnetometers. Sensors, 20(3):753, 2020.

[27] Nikolay Koshev, Anna Butorina, Ekaterina Skidchenko, Alexey Kuzmichev, Alexei Ossadtchi, Maxim Ostras, Maxim Fedorov, and Petr Vetoshko. Evolution of meg: A first meg-feasible fluxgate magnetometer. Human Brain Mapping, 42(15):4844–4856, 2021.

[28] Bourgeois L. A mixed formulation of quasi-reversibility to solve the cauchy problem for laplace’s equation. Inverse Problems, 21(3):1087–1104, 2005.

[29] Etienne Labyt, Marie-Constance Corsi, William Fourcault, Augustin Palacios Laloy, François Bertrand, François Lenouvel, Gilles Cauffet, Matthieu Le Prado, François Berger, and Sophie Morales. Magnetoencephalography with optically pumped 4 he magnetometers at ambient temperature. IEEE transactions on medical imaging, 38(1):90–98, 2018.

[30] Maik Liebl, Frank Wiekhorst, Dietmar Eberbeck, Patricia Radon, Dirk Gutkelch, Daniel Baumgarten, Uwe Steinhoff, and Lutz Trahms. Magnetorelaxometry procedures for quantitative imaging and characterization of magnetic nanoparticles in biomedical applications. Biomedical Engineering/Biomedizinische Technik, 60(5):427–443, 2015.

[31] ME Limes, EL Foley, TW Kornack, S Caliga, S McBride, A Braun, W Lee, VG Lucivero, and MV Romalis. Portable magnetometry for detection of biomagnetism in ambient environments. Physical Review Applied, 14(1):011002, 2020.

[32] F Ludwig, S Mäuselein, E Heim, and M Schilling. Magnetorelaxometry of magnetic nanoparticles in magnetically unshielded environment utilizing a differential fluxgate arrangement. Review of scientific instruments, 76(10):106102, 2005.

[33] Mikhail Malovichko, Nikolay Koshev, Nikolay Yavich, Alexandra Razorenova, and Maxim Fedorov. Electroencephalographic source reconstruction by the finite-element approximation of the elliptic cauchy problem. IEEE Transactions on Biomedical Engineering, 68(6):1811–1819, 2020.

[34] Roshin C Mathew and Christopher M Kramer. Recent advances in magnetic resonance imaging for peripheral artery disease. Vascular Medicine, 23(2):143–152, 2018.

[35] D.W. McRobbie, E.A. Moore, M.J. Graves, and M.R. Prince. MRI from Picture to Proton. Cambridge University Press, 2003.

[36] V. M. Klibanov, N. A. Koshev, L.I. Jingzhi, and A. G. Yagola. Numerical solution of an ill-posed cauchy problem for a quasilinear parabolic equation using a carleman weight function. Journal of Inverse and Ill-posed Problems, 2016.

[37] Madhavan Nair, Rakesh Guduru, Ping Liang, Jeongmin Hong, Vidya Sagar, and Sakhrat Khizroev. Externally controlled on-demand release of anti-hiv drug using magneto-electric nanoparticles as carriers. Nature communications, 4(1):1–8, 2013.

[38] M. Nisenoff and S. Wolf. Observation of a *cosφ* term in the current-phase relation for ”dayem”-type weak link contained in an rf-biased superconducting quantum interference device. Phys. Rev.B, 12:1712–1714, Sep 1975.

[39] The pandas development team. pandas-dev/pandas: Pandas, February 2020.

[40] Mikhail V Petrenko, Sergei P Dmitriev, Anatoly S Pazgalev, Alex E Ossadtchi, and Anton K Vershovskii. Towards the non-zero field cesium magnetic sensor array for magnetoencephalography. IEEE Sensors Journal, 21(17):18626–18632, 2021.

[41] Alexandra Razorenova, Nikolay Yavich, Mikhail Malovichko, Maxim Fedorov, Nikolay Koshev, and Dmitry V Dylov. Deep learning for non-invasive cortical potential imaging. In Machine Learning in Clinical Neuroimaging and Radiogenomics in Neuro-oncology, pages 45–55. Springer, 2020.

[42] Alexandra Rodzinski, Rakesh Guduru, Ping Liang, Ali Hadjikhani, Tiffanie Stewart, Emmanuel Stimphil, Carolyn Runowicz, Richard Cote, Norman Altman, Ram Datar, et al. Targeted and controlled anticancer drug delivery and release with magnetoelectric nanoparticles. Scientific reports, 6(1):1–14, 2016.

[43] Gian Luca Romani, Samuel J Williamson, and Lloyd Kaufman. Biomagnetic instrumentation. Review of Scientific Instruments, 53(12):1815–1845, 1982.

[44] J. Sarvas. Basic mathematical and electromagnetic concepts of the biomagnetic inverse problem. Physics in Medicine and Biology, 32(1):11–22, 1987.

[45] E. Skidchenko, A. Butorina, M. Ostras, P. Vetoshko, A. Kuzmichev, N. Yavich, M. Malovichko, and N. Koshev. Yttrium-iron garnet magnetometer in meg: Advance towards multi-channel arrays. bioRxiv, 2022.

[46] Emmanuel Stimphil, Abhignyan Nagesetti, Rakesh Guduru, Tiffanie Stewart, Alexandra Rodzinski, Ping Liang, and Sakhrat Khizroev. Physics considerations in targeted anticancer drug delivery by magnetoelectric nanoparticles. Applied Physics Reviews, 4(2):021101, 2017.

[47] Tim M Tierney, Stephanie Mellor, George C O’Neill, Niall Holmes, Elena Boto, Gillian Roberts, Ryan M Hill, James Leggett, Richard Bowtell, Matthew J Brookes, et al. Pragmatic spatial sampling for wearable meg arrays. Scientific reports, 10(1):1–11, 2020.

[48] Peter M Vetoshko, Vadim B Volkovoy, Vladimir N Zalogin, and Andrey Yu Toporov. Measuring low alternating magnetic fields by means of bi-containing rare-earth ferrite-garnet films with planar anisotropy. Journal of applied physics, 70(10):6298–6300, 1991.

[49] Pauli Virtanen, Ralf Gommers, Travis E. Oliphant, Matt Haberland, Tyler Reddy, David Cournapeau, Evgeni Burovski, Pearu Peterson, Warren Weckesser, Jonathan Bright, Stéfan J. van der Walt, Matthew Brett, Joshua Wilson, K. Jarrod Millman, Nikolay Mayorov, Andrew R. J. Nelson, Eric Jones, Robert Kern, Eric Larson, C J Carey, Ilhan Polat, Yu Feng, Eric W. Moore, Jake VanderPlas, Denis Laxalde, Josef Perktold, Robert Cimrman, Ian Henriksen, E. A. Quintero, Charles R. Harris, Anne M. Archibald, Antônio H. Ribeiro, Fabian Pedregosa, Paul van Mulbregt, and SciPy 1.0 Contributors. SciPy 1.0: Fundamental Algorithms for Scientific Computing in Python. Nature Methods, 17:261–272, 2020.

[50] Xue Yang, Guoqing Shao, Yanyan Zhang, Wei Wang, Yu Qi, Shuai Han, and Hongjun Li. Applications of magnetic particle imaging in biomedicine: Advancements and prospects. Frontiers in physiology, page 1093, 2022.

[51] N Yavich, N Koshev, M Malovichko, Alexandra Razorenova, and M Fedorov. Conservative finite element modeling of eeg and meg on unstructured grids. IEEE Transactions on Medical Imaging, 41(3):647–656, 2021.

